# Whole genome sequencing enables definitive diagnosis of Cystic Fibrosis and Primary Ciliary Dyskinesia

**DOI:** 10.1101/438838

**Authors:** Jamie M Ellingford, Glenda Beaman, Kevin Webb, Christopher O’Callaghan, Robert A Hirst, on behalf of the 100,000 Genomes Project, Graeme CM Black, William G Newman

**Affiliations:** North West Genomic Laboratory Hub, Manchester Centre for Genomic Medicine, Manchester University Hospitals NHS Foundation Trust, St Mary’s Hospital, Manchester, UK; Division of Evolution and Genomic Sciences, Neuroscience and Mental Health Domain, School of Biological Sciences, Faculty of Biology, Medicine and Health, University of Manchester, Manchester, UK; Manchester Adult Cystic Fibrosis Centre, Manchester University Hospitals NHS Foundation Trust, Manchester, UK; Respiratory, Critical Care and Anaesthesia, UCL Great Ormond Street Institute of Child Health & Great Ormond Street Children’s Hospital & NIHR Great Ormond Street Hospital Biomedical Research Centre, London, UK; Centre for PCD Diagnosis and Research, Department of Infection, Immunity and Inflammation, RKCSB, University of Leicester, Leicester, UK

## Abstract

Understanding the genomic basis of inherited respiratory disorders can assist in the clinical management of individuals with these rare disorders. We apply whole genome sequencing for the discovery of disease-causing variants in the non-coding regions of known disease genes for two individuals with inherited respiratory disorders. We describe analysis strategies to pinpoint candidate non-coding variants within the non-coding genome and demonstrate aberrant RNA splicing as a result of deep intronic variants in *DNAH11* and *CFTR*. These findings confirm clinical diagnoses of primary ciliary dyskinesia and cystic fibrosis, respectively.

## Introduction

Understanding the genomic basis of inherited respiratory disorders, including cystic fibrosis, can assist in the clinical management of individuals with these rare disorders and facilitate testing of at risk relatives. As a result, a number of assays have been developed and adopted in clinical practice to identify disease-causing variation.[1] Genetic testing is undertaken in a staged manner when a diagnosis of cystic fibrosis is suspected. Initial testing involves the genotyping of a number of known disease-causing variants within a specified ethnic group, which can account for up to 95% of molecular diagnoses.[2] When this initial testing does not confirm a diagnosis of cystic fibrosis these tests are supplemented by DNA sequencing techniques to survey the protein-coding regions of genes known as a cause of inherited respiratory disease and related syndromic disorders (Table 1).[3]

**Table 1.** Step-wise genomic assessments for individuals with inherited respiratory disorders.

| **Testing Strategy:** | **Appropriate technologies:** | **Testing for Proband 1:** | **Testing for Proband 2:** |
| --- | --- | --- | --- |
| 1. Assess established disease-causing mutations | Genotyping arrays<br>Direct mutation screening | Direct mutation assessment, 14 mutations (Dec 1996)<br>***CFTR p.Phe508del het*** | Elucigene, CF-EU2 kit (Mar 2011)<br>***negative*** |
| 2. Discover disease-causing mutations in protein coding regions of known disease genes | Gene panel DNA sequencing<br>WES (with virtual gene panels) | MRC-Holland MLPA P091 (Mar 2013)<br>***negative***<br>CFTR exon sequencing (Mar 2013)<br>***CFTR p.Phe508del het*** | Invitae, 31 gene panel (Mar 2016)<br>***DNAH11 p.Tyr2870Ter het*** |
| 3. Discover disease-causing mutations in non-coding and regulatory regions of known disease genes | WGS (with virtual gene panels) | UK 100,000 genomes project (Aug 2018)<br>***CFTR p.Phe508del het***<br>***CFTR c. 3874-4522 A>G het*** | UK 100,000 genomes project (Aug 2018)<br>***DNAH11 p.Tyr2870Ter het***<br>***DNAH11 c.6547-963 G>A het*** |
| 4. Novel disease gene discovery | WES or WGS | n/a | n/a |

WES, whole exome sequencing; WGS, whole genome sequencing; MLPA, multiplex ligation probe amplification; *het*, heterozygous

For individuals with inherited respiratory disorders, whole genome sequencing provides an opportunity to determine the contribution of variation within the non-coding genome in the pathogenesis of well-studied monogenic and heterogeneous disorders, e.g. cystic fibrosis and primary ciliary dyskinesia. These datasets are becoming readily available with the decreasing costs of DNA sequencing. However, establishing routine methods to accurately pinpoint disease-causing variation within large and complex genomic datasets is a significant complication. In this study we describe a framework for the identification of disease-causing variation in the non-coding genome from WGS. We demonstrate the clinical utility of these techniques for individuals without a genomic diagnosis from established diagnostic testing pathways for inherited respiratory disorders. Both our described cases were recruited to the 100,000 Genomes Project with single heterozygous pathogenic variants identified in an autosomal recessive disease gene through previous genetic testing (Table 1).

## Methods

### Genetic testing strategies prior to whole genome analysis

Both probands were assessed systematically for disease-causing mutations in inherited respiratory disorder genes, as outlined in Table 1.

### Whole genome sequencing analysis strategies

Whole genome sequencing datasets were created through the UK 100,000 genomes project main program, using Illumina X10 sequencing chemistry. Sequencing reads were aligned to build GRCh37 of the human reference genome utilizing Issac (Aligner). Small variants were identified through Starline (SNV and small indels ≤ 50bp), and structural variants were identified utilizing Manta and Canvas (CNV Caller). Variants were annotated and analysed with the Ensembl variant effect predictor (v92) and bespoke perl scripts within the Genomics England secure research embassy. Prioritized variants were: (i) rare (<0.01%) or absent in gnomAD (available at http://gnomad.broadinstitute.org), (ii) rare (<1%) or absent within the Genomics England Pilot cohort, and (iii) within a defined genomic locus. We defined the relevant genomic loci as: *CFTR* (chr7:117,120,017-117,308,718, *GRCh37*) for proband 1, and *DNAH11* (chr7:21,582,833-21,941,457, *GRCh37*) for proband 2.

### RNA investigations

Lymphoblast cell cultures were established for a control sample and proband 1. RNA was extracted using the RNeasy^®^ Mini Kit (Qiagen, UK, Catalogue No. 74104) following the manufacturer’s protocol. RNA was extracted from whole-cell blood using the PAXgene™ Blood RNA System Kit (Qiagen, UK. Catalogue No. 762174), following the manufacturer’s protocol for a control sample and proband 2. Extracted RNA was reverse transcribed using the High Capacity RNA to cDNA Kit (Applied Biosystems, UK. Catalogue No. 4387406) following the manufacturer’s protocol. Gene specific primers (available on request) amplified relevant regions of *CFTR* and *DNAH11*. PCR products were visualized on an agarose gel using a BioRad Universal Hood II and the Agilent 2200 Tapestation. Visualized bands were cut out and prepared for capillary sequencing on an ABI 3730×l DNA Analyzer.

### Variant Segregation

Primers were designed to amplify relevant genomic DNA regions in the probands and their unaffected relatives (both parents for proband 2 and daughter for proband 1). Amplicons were prepared for capillary sequencing using an ABI 3730×l DNA Analyzer.

## Results

### Clinical findings

Proband 1 (Figure 1) received a late diagnosis of cystic fibrosis at 17 years old. Genetic testing described in Table 1 uncovered the common del508 mutation in *CFTR* in a heterozygous state. A sweat test was positive: sweat chloride 68 mmol/L (normal range = 0-39 mmol/L); sweat conductivity 92 mmol/L (normal range = 0-49 mmol/L). The proband is currently 51 years old but her disease severity has progressed and she is awaiting double lung transplant.

**Figure 1.**
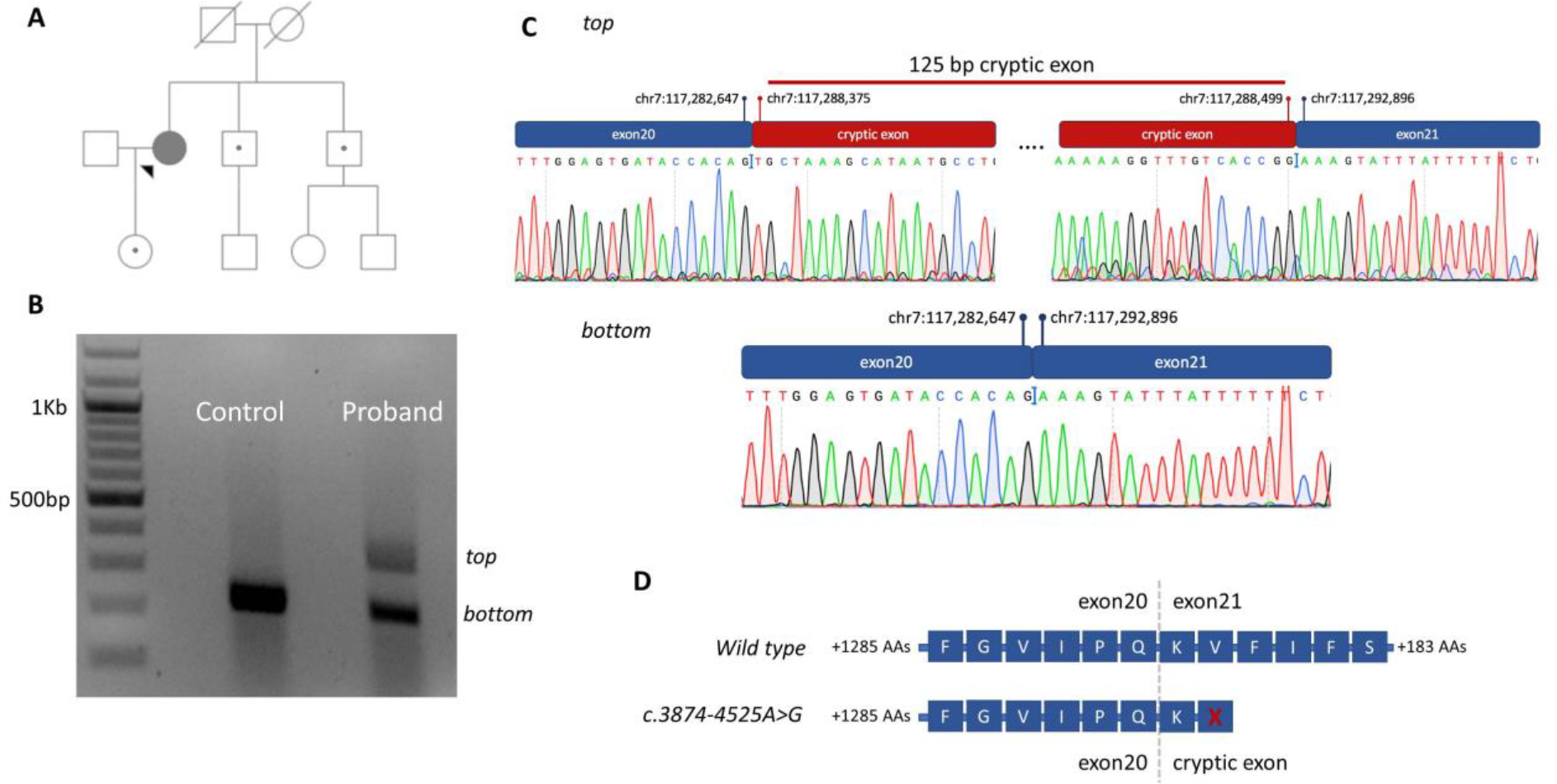
*CFTR* c.3874-4525A>G identified in proband 1. **(A)** Family pedigree showing the proband and unaffected relatives. Carriers of *CFTR* p. Phe508del are indicated with closed circles within an open symbol. **(B)** Gel electrophoresis results for the proband and control sample, visualized using a BioRad Universal Hood II. RNA was reverse transcribed after extraction from lymphoblast cell lines and then amplified using primers specific to exons 20 and 21 of the *CFTR* gene (NM_000492.3). The caption shows two distinct cDNA amplicons in the proband sample separated by ~100 base pairs. **(C)** Sanger sequencing chromatograms for the *CFTR* gene generated after cutting out relevant cDNA PCR products from the agarose gel. The larger (*top*) cDNA PCR product shows a 125 base pair cryptic exon as a result of c.3874-4525A>G. **(D)**Impact of the cryptic exon on the translated protein. A stop codon is expected to be introduced within the cryptic exon and will result in premature termination of protein synthesis. Amino acids (AAs) are provided with single letter notations, with *X* indicating a stop codon. Vertical intersects indicate transition of the cDNA to the adjacent exon.

Proband 2 (Figure 2) was diagnosed in childhood with bronchiectasis, she is currently 54 years of age. Nasal nitric oxide levels were extremely low at 4 parts per billion (*ppb*, normal range = <25 ppb) consistent with a diagnosis of PCD.[4] Three examinations of the proband’s cilia with electron microscopy (EM) showed a significant proportion of static and dyskinetic cilia (Supplementary Video), with a high ciliary beat frequency: 20.2Hz (95%CI=19.8-20.5Hz); 22.0Hz (95%CI=20.1-23.2Hz); and 20.1Hz (95%CI=19.8–20.5Hz). EM histology showed normal dynein arms and microtubules with no ciliary disorientation, and conical ciliated protrusions were observed from epithelial cells. These findings are consistent with mutations in *DNAH11*.[5] Genetic testing described in Table 1 identified a heterozygous nonsense mutation in *DNAH11*.

**Figure 2.**
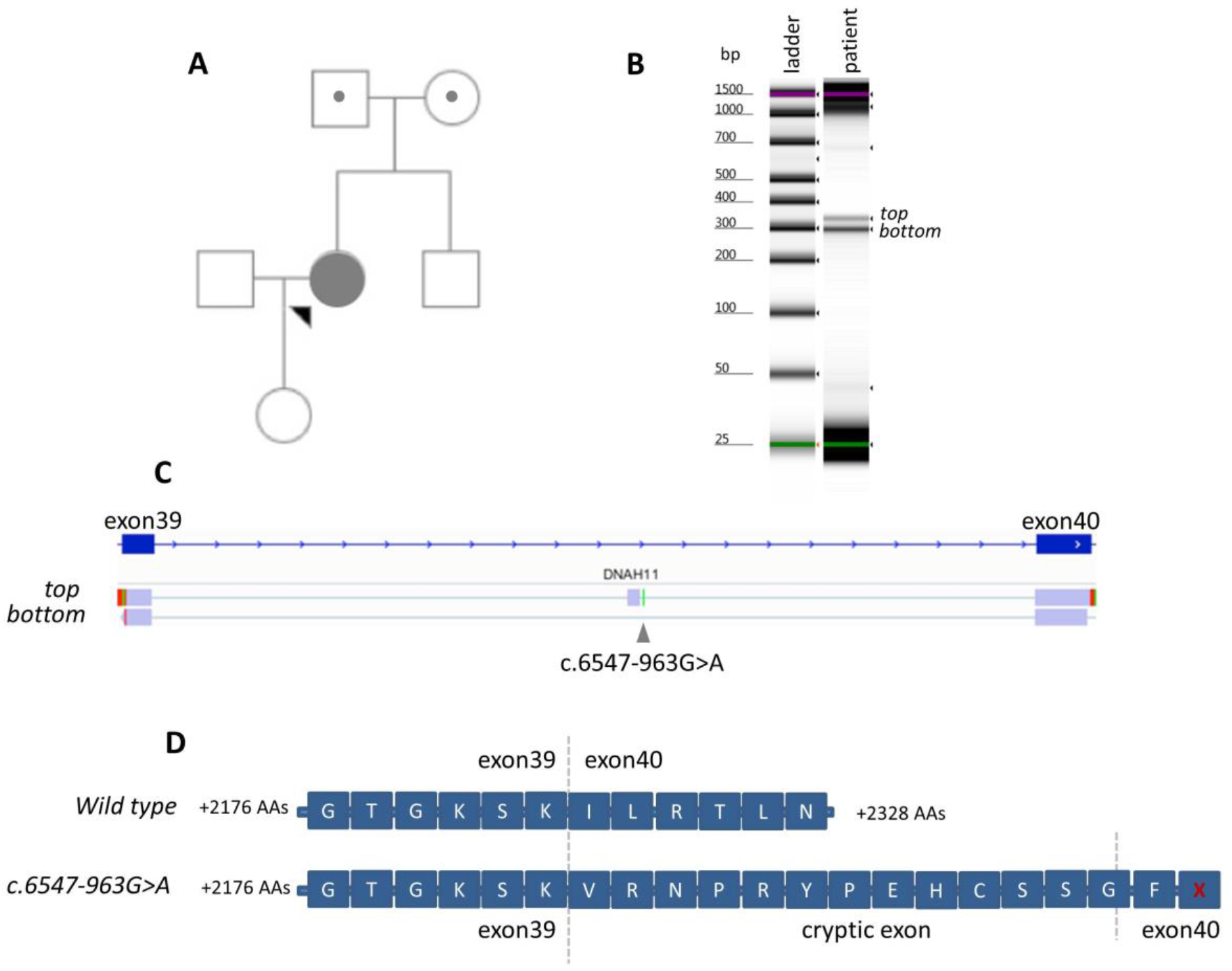
*DNA11* c.6547-963 G>A identified in proband 2. **(A)** Family pedigree showing the proband and her unaffected father and mother who carry heterozygous alleles of *DNAH11* c.8610C>G and c.6547-963G>A, respectively. **(B)** Gel electrophoresis results for the proband, visualized using an Agilent 2200 Tapestation. RNA was reverse transcribed after extraction from whole blood and then amplified using primers specific to exons 39 and 40 of the *DNAH11* gene (NM_001277115.1). The caption shows two distinct cDNA amplicons in the proband sample separated by ~40 base pairs. **(C)** Integrated Genomic Viewer snapshot of the alignment of sequencing products to the human reference genome (GRCh37) showing the introduction of a 38 base pair cryptic exon (chr7:21,746,318-21,746,355) as a result of c.6547-963 G>A. The *top* and *bottom* bands were sequenced after being cut from an agarose gel electrophoresis. **(D)** Impact of the cryptic exon on the translated protein. The cryptic exon shifts the reading frame and is expected to introduce a premature stop codon in exon 40, resulting in premature termination of protein synthesis. Amino acids (AAs) are provided with single letter notations, with *X* indicating a stop codon. Vertical intersects indicate transition of the cDNA to the adjacent exon.

### Identification of non-coding likely pathogenic variation from WGS

Whole genome sequencing generated 49,764 unique reads for the specified region of *CFTR*, achieving coverage of 30x and 10x for 91.5% and 99.3%, respectively. Similarly, 92,053 unique reads were generated for the specified region of *DNAH11*, achieving coverage of 30x and 10x for 89.8% and 100%, respectively. Our variant analysis strategy for WGS identified both previously identified pathogenic variants (Table 1). In addition, these analyses identified candidate heterozygous variants within the introns of *CFTR* and *DNAH11* (Table 1).

#### DNAH11 c.6547-963 G>A

This variant has not been previously reported as a cause of disease. The variant is present as two heterozygous alleles in gnomAD (rs764374746). The variant alters the nearby sequence motif from <u>G</u>GGT to <u>A</u>GGT. Analysis with the *in-silico* splicing tools SpliceSiteFinder-like, MaxEntScan and Human Splicing Finder suggest that this could result in a new cryptic splice donor within this sequence motif.

#### CFTR c. 3874-4522 A>G

This variant has previously been reported in a single case of cystic fibrosis.[6] The variant is absent from gnomAD. *In-silico* splicing tools (SpliceSiteFinder-like, MaxEntScan and Human Splicing Finder) suggest that this variant may result in the introduction of a new cryptic splice acceptor site. The variant is not included in diagnostic genotyping kits and is not surveyed through protein-coding based DNA sequencing techniques.

### Variant Segregation

Segregation analyses identified that *DNAH11* c.8610C>G and c.6547-963G>A were *in-trans* in proband 2, who had inherited these variants from her unaffected father and mother, respectively (Figure 1).

Assessment of the unaffected daughter of proband 1, who had inherited p. Phe508del on a single allele, also demonstrated that *CFTR* c.3874-4522 A>G and p.Phe508del are *in-trans* (Figure 1).

### RNA assessments for variant pathogenicity

PCR products were amplified from proband and control cDNA utilizing primers designed for exons flanking the prioritized intronic variants. Our findings show abnormal RNA splicing patterns for the probands carrying *DNAH11* c.6547-963 G>A and *CFTR* c. 3874-4522 A>G through the introduction of cryptic splice sites (Figures 1 & 2). Both cryptic exons are expected to result in premature termination of protein synthesis and are interpreted as pathogenic (ClinVar submission in process).

## Discussion

In this study we describe the discovery of non-coding pathogenic variants from WGS as a cause of inherited respiratory disorders in two unrelated individuals. Through targeted RNA investigations we established that these prioritized variants have adverse effects on RNA expression (Figures 1 & 2). These findings in *CFTR* and *DNAH11* confirm clinical diagnoses of autosomal recessive cystic fibrosis and primary ciliary dyskinesia, respectively, and enable informed management and cascade testing for at risk relatives. The delineation of the underlying mutations in *CFTR* is an important consideration for the lifelong management for individuals with cystic fibrosis, as appropriate modulator therapies can extend life span.[7] We pinpoint mutations that directly impact RNA splicing for *CFTR* and *DNAH11*, and thereby underline the potential utility of antisense oligonucleotide therapies which target and correct aberrant splicing within pre-mRNA.[8] Both patients are in their sixth decade of life and present symptoms consistent with a milder spectrum of disease. The hypothesis that alternative splicing mechanisms retain some residual normal transcript, and thereby underpin milder phenotypic presentation, provides an interesting avenue for future investigations.

Our findings demonstrate the additional advantage of WGS over other commonly used genetic testing modalities (Table 1). Other technologies could have uncovered the pathogenic variants discovered in this study. For example, assessment of RNA expression through mRNA sequencing [9] or targeted sequencing of intronic regions of *CFTR* and *DNAH11*. However, in addition to the discovery of novel non-coding pathogenic variation, WGS also provided a one-step tool to test and explicitly rule out other hypotheses that may be missed by these other techniques, e.g. the existence of novel structural variants and variants impacting the promoter regions of *CFTR* and *DNAH11*. Moreover, if patients are negative from these hypothesis-led assessments, then WGS datasets can lead to the discovery of new disease-causing genes (Table 1).[3] Our analysis strategy was guided by the clinical presentation of the recruited probands and the existence of heterozygous pathogenic variants in genes already established as a cause of inherited respiratory disorders. We demonstrate here, and previously,[10] that such strategies can be rapidly and effectively applied to WGS datasets for families with significant previous genomic testing to uncover novel genomic diagnoses. We suggest that these strategies are applied to individuals whose clinical presentation is in keeping with an autosomal recessive Mendelian disorder and previous testing of known disease genes has identified a single heterozygous pathogenic variant.

## Acknowledgements

J.M.E is funded by a postdoctoral research fellowship from the Health Education England Genomics Education Programme (HEE GEP). W.G.N is supported through the Manchester NIHR BRC. The views expressed in this publication are those of the authors and not necessarily those of the HEE GEP. This research was made possible through access to the data and findings generated by the 100,000 Genomes Project. The 100,000 Genomes Project is managed by Genomics England Limited (a wholly owned company of the Department of Health). The 100,000 Genomes Project is funded by the National Institute for Health Research and NHS England. The Wellcome Trust, Cancer Research UK and the Medical Research Council have also funded research infrastructure. The 100,000 Genomes Project uses data provided by patients and collected by the National Health Service as part of their care and support.

